# Ontologizer 3: a cross-platform desktop application for frequentist and Bayesian GO enrichment analysis

**DOI:** 10.64898/2026.06.04.730045

**Authors:** Lukas Ramlow, Jasmin Scholtes, Daniel Danis, Peter N. Robinson

## Abstract

We present Ontologizer 3, an easy-to-use cross-platform desktop application for Gene Ontology (GO) overrepresentation analysis. Ontologizer 3 offers two complementary methods. The first is a frequentist approach that evaluates GO terms individually using a one-sided Fisher’s exact test, yielding term-level significance values. The second is a Bayesian approach that jointly evaluates terms using model-based gene set analysis (MGSA), yielding term-level posterior probabilities. Due to substantial overlap among annotated gene sets resulting from the GO’s hierarchical structure, the two methods produce different results. The frequentist approach tends to report a lengthy list of terms with strong annotation overlap, whereas, MGSA yields a parsimonious set of terms that explain the observed gene activity. Using simulated data with a known ground truth, we demonstrate that both methods reliably identify the causal term, but MGSA achieves substantially higher precision. Ontologizer 3 is implemented as a Tauri application with a Rust backend and an Angular frontend, presenting enriched terms in tabular and graphical form. The software is freely available under the MIT licence at https://github.com/P2GX/ontologizer-gui. Installation packages for Macintosh, Windows and Debian-based Linux are available on the GitHub Releases page.

## Introduction

The Gene Ontology (GO) [3, 1] provides terms to describe functions of gene products, i.e., proteins encoded by genes or non-coding RNA molecules. GO terms are interconnected by relations that link the nodes in the graph. The most important of these are `‘‘is-a’’`, which denotes a subclass relationship, and `‘‘part-of’’`, which denotes a part-whole relationship (e.g. ‘the nucleus is part of a cell’). Genes are linked to GO terms by means of GO annotations, which are statements that describe attributes of genes in three categories (arranged as the three subontologies of GO). *Molecular Function* describes the normal molecular activity of a gene product; for instance *DNA helicase activity* (GO:0003678) is defined as the “Unwinding of a DNA helix, driven by ATP hydrolysis” and is used to annotate gene products including Chromodomain helicase DNA binding protein 1 like (*CHD1L*). *Biological Process* describes the pathways and larger processes to which the gene product’s activity contributes. For instance, Collagen alpha-2(XI) chain (*COL11A2*) is annotated to *cartilage development* (GO:0051216). Finally, *Cellular Component* denotes where the gene product is located when the activity occurs. *Golgi membrane* (GO:0000139) annotates Galactosylgalactosylxylosylprotein 3-beta-glucuronosyltransferase (*B3GAT2*).

By the transitivity principle or “true-path rule”, a positive annotation to a GO term implies annotation to all its “is-a” and “part-of” parents and ancestors. For instance, this implies that *COL11A2* is also annotated to *skeletal system development* (GO:0001501) because cartilage is part of the skeletal system.

One of the most important applications of GO is in the analysis of lists of differentially expressed genes derived from exploratory experiments in which the transcriptional activity of all or most genes is assayed with RNA-seq or comparable methods. A large number of statistical frameworks have been developed to identify enriched biological categories in such gene lists [17, 11], spanning both frequentist [8, 22, 6] and Bayesian [23, 16, 15] approaches. One such method is GO overrepresentation analysis (ORA), which asks whether more genes annotated to a given GO term are found to be differentially expressed than one would expect by chance If so, then we say that genes annotated to the GO term are “overrepresented” in the set of differentially expressed genes [20].

The standard ORA approach involves performing a Fisher’s exact test (FET) for each GO term separately [8, 19]. For this reason, we refer to the standard approach as “term-for-term” (TfT) [4]. A direct consequence of the transitivity principle is that terms in ancestor-descendant relations demonstrate a high degree of overlap of their annotated genes. This implies that TfT tends to return large numbers of correlated categories, leaving the choice of the most relevant ones to the user’s interpretation [2, 13]. To address this issue, the model-based gene set analysis (MGSA) evaluates all categories at once by embedding them in a Bayesian network, in which gene response is modeled as a function of the activation of biological categories. Probabilistic inference is used to identify the active categories. This Bayesian modeling approach naturally takes term overlap into account and avoids the need for multiple-testing correction by jointly inferring posterior probabilities for all terms [5].

We previously presented versions 1 and 2 of the Ontologizer as desktop and web applications in Java 1.1 and 1.4 [21, 4]. These applications can no longer be run with modern Java versions. Here we present Ontologizer 3, implemented in Rust with an Angular front-end packaged via the Tauri framework as a cross-platform native desktop application. Ontologizer 3 implements both TfT (with the standard suite of multiple-testing corrections) and an improved MGSA. As with previous versions, our goal is to provide an easy-to-use desktop application for running GO enrichment analysis and visualizing the results in tabular and graphical form.

## Availability and Workflow

Ontologizer 3 is implemented in Rust with an Angular front-end, packaged as a cross-platform native desktop application via the Tauri framework. Pre-built installers for Windows (.msi, .exe), macOS (.dmg), and Linux (.AppImage, .deb, .rpm) are distributed via the GitHub Releases page. Ontologizer 3 is freely available under the MIT license. The relevant repositories and resources are:

- Frontend: https://github.com/P2GX/ontologizer-gui
- Backend: https://github.com/P2GX/ontologizer
- Documentation: https://p2gx.github.io/ontologizer-gui/

The analysis proceeds in three steps (Figure 1). First, Ontologizer requires the Gene Ontology in JSON format, an organism-specific GO association file (GAF), and two plain-text files listing the population and study genes with one gene symbol per line. The Gene Ontology and GAF files for commonly studied organisms can be downloaded from within Ontologizer 3, whereas the population and study gene files are experiment-specific and must be provided locally. Gene symbols in these files must match the *DB Object Symbol* (column 3) of the GAF file. Second, the user selects either “Statistical Testing”, to run TfT with a chosen multiple-testing correction, or “Bayesian Inference”, to run MGSA (see Methods for details).

**Figure 1.**
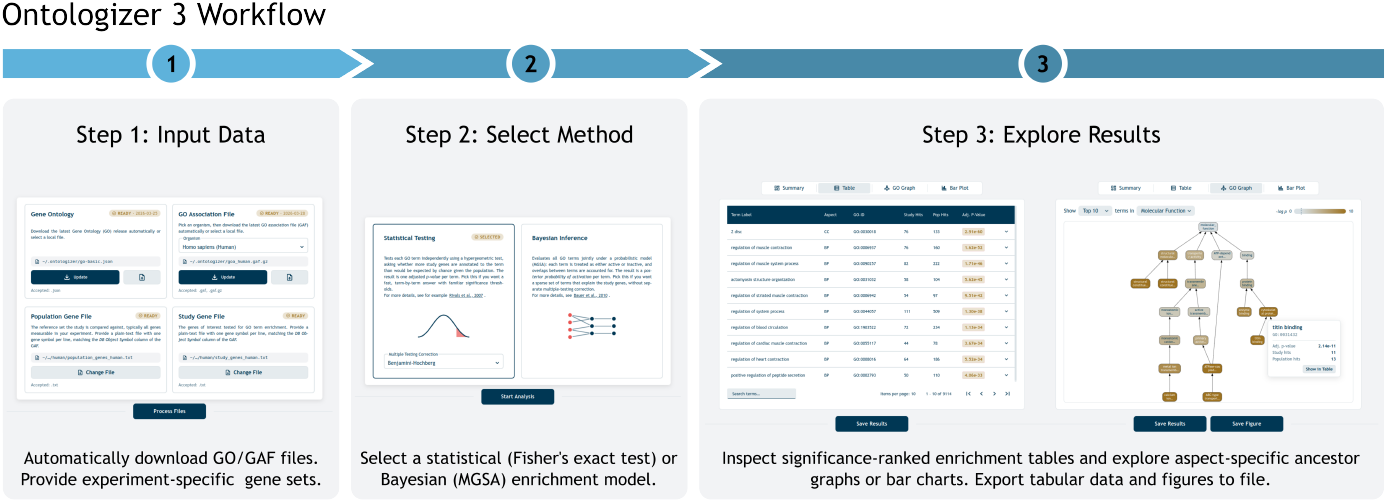
Ontologizer 3 Workflow: Step 1: Provide the GO and organism-specific GAF files, as well as the population and study gene lists specific to the experiment. GO and GAF files can be downloaded within the application. Step 2: Select either Fisher’s exact test with a multiple testing correction method (statistical testing) or model-based gene set analysis (Bayesian inference) for term enrichment analysis. Step 3: Explore the enriched terms in a ranked, searchable table, an interactive ancestor graph filtered by GO aspect, and a bar plot. Both the tabular results and the figures can be exported.

Third, enriched terms are presented in a searchable table, ranked by adjusted *p*-value or posterior probability depending on the method. The results can additionally be explored as an interactive ancestor subgraph of the Gene Ontology, filtered by GO aspect and constructed from either all significant terms or the top 10, 25, or 50 ranked terms, or as a bar chart of ranked terms. The bar chart can display the significance measure itself, the number of study-set genes annotated to each term, or the log-fold change in the number of annotated genes between the study and population sets. Tabular results and figures can be exported as CSV and PNG files, respectively.

## Methods

Ontologizer 3 provides a statistical testing and a Bayesian inference approach to assess enrichment of GO terms. The two methods are detailed in the following.

### Term-for-term

TfT uses Fisher’s exact test to evaluate term overrepresentation [19]. Let *N* denote the total number of genes in the population set, and *n* the number of genes in the study set. For a given term *T*, let *K* be the number of genes in the population set annotated to *T*, and *k* the number of genes in the study set annotated to *T*. The probability of observing exactly *k* such *NAR Genomics and Bioinformatics, YEAR, Volume XX, Issue x* genes follows the hypergeometric distribution

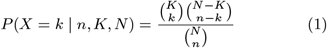

and the one-sided *p*-value for over-representation is the right tail

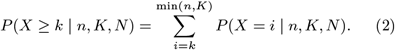

### Multiple Testing Correction

Because a typical analysis tests thousands of GO terms *T*_1_, …, *T*_*m*_, Ontologizer adjusts the raw *p*-values from Eq. 2 for multiple testing [18]. Specifically, we test every term that has at least one gene from the study set annotated. Let *p*_(1)_≤ … ≤ *p*_(*m*)_ denote the sorted *p*-values of the *m* tested terms. Four options are provided: ‘None’ 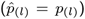, ‘Bonferroni’ 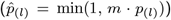, and ‘Bonferroni-Holm’ 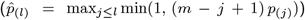 [14] to control the family-wise error rate and ‘Benjamini-Hochberg’ 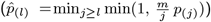 [7] to control the false-discovery rate.In all cases Ontologizer reports the adjusted *p*-value 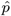for the chosen procedure.

### Model-based gene set analysis

The Bayesian approach uses MGSA to determine the posterior probability of term activation. MGSA differs from TfT in two key ways. First, rather than associating term activation with the rejection of a null hypothesis, it specifies a generative model and directly computes the posterior probability of term activation given the observations [12]. Second, MGSA evaluates all terms simultaneously, aiming to identify a parsimonious set of terms that together explain the observed gene activity. The algorithm has been thoroughly described in [5]; here we review the key concepts and improvements implemented in the current version.

### Model

MGSA models gene response using a three-layer Bayesian network of terms (*T*), hidden gene states (*H*), and observed gene states (*O*) (Figure 2). The term layer consists of *m* Boolean nodes, one for each GO term. The hidden and observed layers each consist of *n* Boolean nodes, one for each gene. Edges from the term layer to the hidden layer are determined by the GO associations: a term is connected to every gene annotated to it under the true-path rule. Edges from the hidden to the observed gene layer are one-to-one. The objective of MGSA is to determine the posterior probability of term activation, *P* (*T*_*i*_= 1|*O*). To this end, we need to establish how term states propagate to the observed gene layer. MGSA assumes that each term is independently active with a prior probability *p*. The hidden layer represents the true, unobserved activation of the genes. We assume that every gene connected to at least one active term is active itself. The observed gene layer represents the experimentally measured activation of the genes and is a noisy realization of the hidden layer. This noise is parameterized by two hyperparameters, *α* and *β*, where *α* is the probability that a gene that is inactive in the hidden layer is observed as active, and *β* is the probability that a gene that is active in the hidden layer is observed as inactive. Together, these assumptions allow MGSA to infer which terms most likely explain the observed gene activity.

**Figure 2.**
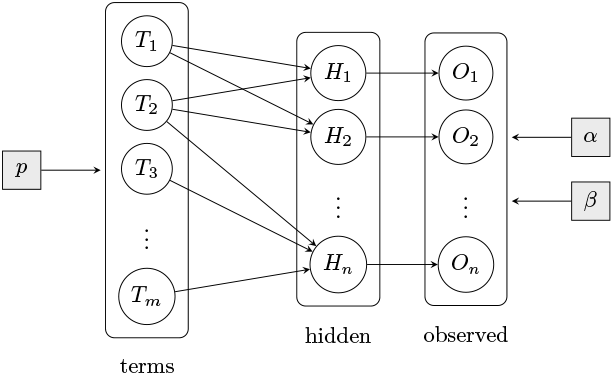
Bayesian network underlying MGSA. Terms *T*_1_, …, *T*_*m*_ are activated independently with prior probability *p*. An active term activates all genes annotated to it in the hidden layer *H*_1_, …, *H*_*n*_. The observed states *O*_1_, …, *O*_*n*_ are noisy realizations of the hidden states, governed by the false-positive rate *α* and the false-negative rate *β*. Hyperparameters are drawn next to the layer they affect.

More formally, calculating the marginal posterior probability of activation, *P* (*T*_*i*_= 1|*O*), for each term *T*_*i*_requires the joint posterior, which by Bayes’ rule is:

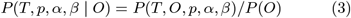

where *P* (*T, O, p, α, β*) is obtained from the joint distribution of the hierarchical model

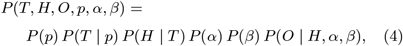

by marginalizing over the hidden layer *H*. In order to evaluate eq. (3), we must specify the conditional probability distributions (CPDs) of each factor in eq. (4). First, we consider the CPDs that remain unchanged from the original algorithm [5], before discussing the priors *P* (*p*), *P* (*α*) and *P* (*β*), which constitute the main methodological change in the present implementation. The prior activation of each term *T*_*i*_*\* T* is described by a Bernoulli distribution *P* (*T*_*i*_= 1 | *p*) = *p* so that the probability for a specific configuration *T* with *m*_1_active and *m*_0_inactive terms is given by

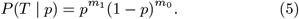

The hidden and observed states factorize across genes

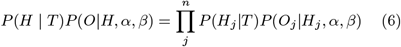

because each *H*_*j*_depends only on the terms annotated by gene *j* and each *O*_*j*_depends only on *H*_*j*_. The propagation of activation from the term layer to the hidden layer is deterministic

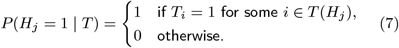

where *T* (*H*_*j*_) is the set of terms annotating gene *j*, i.e. a hidden gene state is active (*H*_*j*_= 1) if at least one term annotated to the gene *j* is active. The observation layer is in turn a noisy realization of the hidden layer:

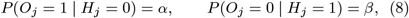

where *α* is the probability that a gene that is inactive in the hidden layer is observed as active (false positive) and *β* is the probability that a gene that is active in the hidden layer is observed as inactive (false negative). Together eqs. (6), (7), and (8) allow the marginalized density to be calculated:

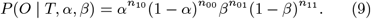

where *n*_*xy*_is the number of genes with observed state *x* and hidden state *y* under the configuration of *T*. This completes the specification of the likelihood *P* (*O* | *T, α, β*) in eq. (3). What remains are the priors *P* (*p*), *P* (*α*), *P* (*β*) which we discuss next.

In the original MGSA implementation, the three hyper-parameters were restricted to a small set of discrete values, primarily due to computational constraints. Here, we use the entire continuous state space together with beta priors, parameterized by a mean *µ* and scale factor *s*. To avoid conflicts with the parameters *α, β* of the model, we depart from the standard notation and denote the Beta shape parameters by

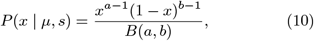

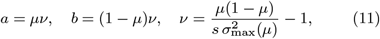

where *x * {p, α, β}* denotes the hyperparameters, *B*(*a, b*) is the beta function and 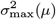 is the largest variance for which the distribution remains unimodal (*a, b >*1):

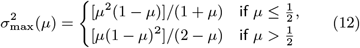

For all three priors we use a scale factor *s* = 0.5 and prior means *µ*_*p*_, *µ*_*α*_, *µ*_*β*_derived empirically from the input data. We first approximate a minimal term set required to cover the active genes using the greedy algorithm of Chvatal [10]. We denote the resulting term configuration *T* ^*\**^. The prior mean for the term-activation probability is the resulting fraction of active terms, *µ*_*p*_= *m*_1_*/m*, where *m*_1_is the number of active terms in *T* ^*\**^and *m* the total number of GO terms. Ideally we would also estimate the mean of *α* and *β* (*µ*_*α*_and *µ*_*β*_) based on *T* ^*\**^. For *β* this is indeed possible. We set *µ*_*β*_= *n*_01_*/*(*n*_01_+ *n*_11_), the fraction of genes that *T* ^*\**^predicts to be active but are observed as inactive. The same strategy fails for *α* because, by construction, *T* ^*\**^covers all observed-active genes, implying *n*_10_= 0. We instead use the marginal probability that any gene is observed as active as an upper bound on *α* and set *µ*_*α*_= *n*_1_*/n*, where *n*_1_is the number of observed-active genes and *n* is the total number of genes in the population.

### Sampling

We approximate the joint posterior *P* (*T, p, α, β* | *O*) using a Metropolis-Hastings (MH) algorithm [9]. In contrast to the original implementation we also apply Bayesian optimization to the three hyperparameters *p, α, β*. At each iteration the sampler proposes either a new term configuration (in 9/10 cases) or a new value for one of the hyperparameters (in 1/10 cases, corresponding to 1/30 cases for each of the hyperparameters).

A proposed move *θ*^*t*^→ *θ*^*p*^with *θ* = (*T, p, α, β*) is accepted with probability

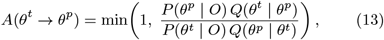

where *Q* is the proposal density.

Sampling over the term configuration *T* follows Bauer et al. [5]. Briefly, a candidate *T* ^*p*^is obtained by either toggling a single term or by swapping the states of one active and one inactive term, and the proposal density *Q*(*T* ^*p*^|*T* ^*t*^) is uniform over the resulting neighbourhood of *T* ^*t*^.

To update a hyperparameter x ∈ * {*p, α, β*} we perturb it in logit space by a zero-mean Gaussian random number *ε \** 𝒩 (0, *σ*) and transform back via the sigmoid function:

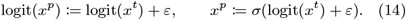

This guarantees that proposed states remain in (0, 1) without rejection or reflection at the boundaries. While the Gaussian step is symmetric in logit space, it is not in the original parameter space. This is captured by the Jacobian that relates the proposal densities in the two spaces: *Q*(*x*^*p*^| *x*^*t*^) = *Q*_*y*_(*y*^*p*^| *y*^*t*^) · |*dy*^*p*^*/dx*^*p*^|. Using |*dy*^*p*^*/dx*^*p*^| = 1*/*[*x*^*p*^(1 *− x*^*p*^)] and the fact that the ratio of the *Q*_*y*_(· | ·) cancels in the acceptance ratio because it is symmetric, the acceptance probability for a hyperparameter update reduces to

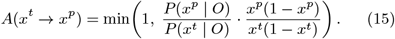

The chain is run for *N*_burn_= 1000 *· m* burn-in iterations, after which *N* = 5000 *· m* samples are drawn. The marginal posterior probability that term *i* is active is estimated samples in which *T*_*i*_= 1,

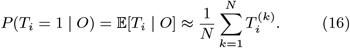

Ontologizer 3 also return estimates of the posterior means of the hyperparameters x ∈ * {*p, α, β*} estimated as the average

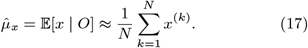

Where *T* ^(*k*)^ and *x*^(*k*)^denote the *k*-th sample after the burn-in period.

### Validation

We validated Ontologizer 3 on a simulated dataset, where the ground truth of active terms is known by construction. This setup allows us to benchmark the precision and recall of the retrieved terms for the provided methods (Figure 3). A Snakemake workflow for reproducibility is available at https://github.com/P2GX/ontologizer-benchmark.

**Figure 3.**
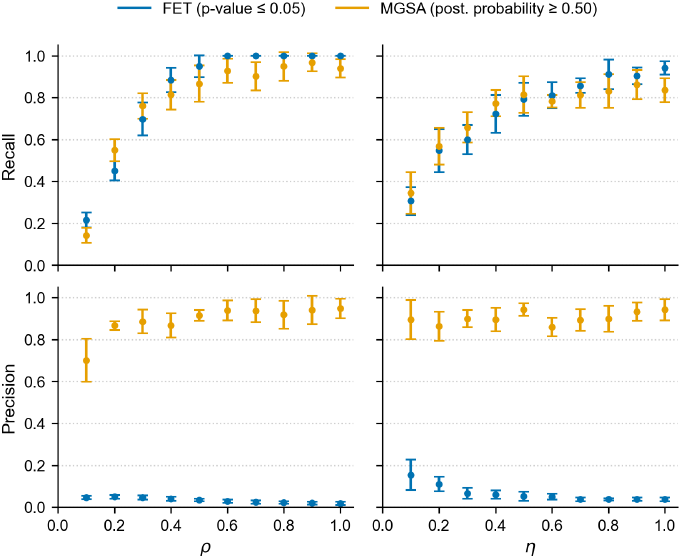
Validation of Ontologizer’s enrichment methods on simulated data. Precision and recall of FET with Bonferroni correction (blue) and MGSA (orange) as functions of *ρ* (the per-term sampling fraction) and *η* (the causal fraction of the study set). When *ρ* is varied, *η* is fixed at 0.5; when *η* is varied, *ρ* is fixed at 0.5. Each point indicates the mean and errorbars the standard deviation over ten simulated study sets per parameter combination.

The simulated population and study sets were constructed as follows. The population set comprises all human protein-coding genes. To construct the study set, we randomly selected GO terms annotated to at least ten genes and declared them *active*. For each active term, a fraction *ρ* of its annotated genes were added to the study set. For example, if *ρ* = 0.5 then 50% of each active term’s annotated genes were added to the study set. This was repeated until the study set contained approximately 500 genes (the exact number depends on the number of genes annotated to the active terms). We then added unrelated noise genes until only a target fraction *η* of the study set was annotated to an active term. For example, if *η* = 0.5 then an equal number of unrelated noise genes and term-related genes have been added to the study set. Each simulated study set is therefore characterized by the list of active GO terms used to construct it, together with the parameters *ρ* and *η*. Because each study set is a random realization, we repeated this procedure ten times for each (*ρ, η*) combination.

We then ran Ontologizer’s TfT with Bonferroni multiple testing correction and MGSA on these dataset and assessed the term recall, i.e. the number of active terms that are also flagged as enriched divided by the number active terms, and the precision, i.e. the number of active terms that are also flagged as enriched divided by the number of terms flagged as enriched. We consider a term to be enriched if its adjusted *p*-value is smaller than a significance threshold of 0.05 or if the posterior probability is larger than 0.5. Figure 2 displays the mean and variance in of term recall and term precision over the ten study sets per parameter combination. We find that both methods achieve high term recall. In other words, the retrieved term lists contain the causative terms in both cases. However, the two algorithms differ strongly in their term precision, which is typically above 80% for MGSA and below 10% for FET.

MGSA therefore not only recovers most of the causal terms but also reports few unrelated ones, whereas FET recovers the causal terms hidden among a large number of redundant ones. As previously discussed, this is because MGSA evaluates terms jointly and naturally accounts for annotation overlap, whereas FET evaluates each term separately.

## Conclusion

We have presented Ontologizer 3, a free, open-source, cross-platform application for Gene Ontology overrepresentation analysis. Ontologizer implements both TfT using the classical Fisher’s exact test with a standard suite of multiple testing corrections, as well as an improved version of MGSA with empirical Beta priors and continuous-value sampling on the hyperparameters. Using simulated data with a known ground truth, we demonstrated that both methods reliably identify enriched terms. However, MGSA returns a parsimonious list of terms, with a precision above 80% compared to a precision of below 10% for the TfT approach. Ontologizer 3 combines these two methods into one easy-to-use application.

## Author contributions statement

L.R.: Methodology, Software, Validation, Data Curation, Writing - Original Draft, Visualization J.S.: Software D.D.: Software P.N.R.: Conceptualization, Methodology, Writing - Review & Editing, Supervision, Funding acquisition

## Acknowledgments

The authors would like to dedicate this work to the memory of Sebastian Bauer, first author of the Ontologizer 2.0 and MGSA papers.

## References

1. S. A. Aleksander, J. P. Balhoff, S. Carbon, J. M. Cherry, D. Ebert, M. Feuermann, P. Gaudet, N. L. Harris, D. P. Hill, P. Kalita, et al. The gene ontology knowledgebase in 2026. Nucleic Acids Research, 54(D1):gkaf1292, 2025.

2. A. Alexa, J. Rahnenführer, and T. Lengauer. Improved scoring of functional groups from gene expression data by decorrelating GO graph structure. Bioinformatics, 22(13): 1600–1607, 2006.

3. M. Ashburner, C. A. Ball, J. A. Blake, D. Botstein, H. Butler, J. M. Cherry, A. P. Davis, K. Dolinski, S. S. Dwight, J. T. Eppig, et al. Gene ontology: tool for the unification of biology. Nature genetics, 25(1):25–29, 2000.

4. S. Bauer, S. Grossmann, M. Vingron, and P. N. Robinson. Ontologizer 2.0—a multifunctional tool for GO term enrichment analysis and data exploration. Bioinformatics, 24(14):1650–1651, 2008.

5. S. Bauer, J. Gagneur, and P. N. Robinson. GOing Bayesian: Model-based gene set analysis of genome-scale data. Nucleic Acids Research, 38(11):3523–3532, 2010.

6. M. Bayerlová, K. Jung, F. Kramer, F. Klemm Bleckmann, and T. Beißbarth. Comparative study on gene set and pathway topology-based enrichment methods. BMC Bioinformatics, 16(1):334, 2015.

7. Y. Benjamini and Y. Hochberg. Controlling the False Discovery Rate: A Practical and Powerful Approach to Multiple Testing. Journal of the Royal Statistical Society Series B: Statistical Methodology, 57(1):289–300, 1995.

8. E. I. Boyle, S. Weng, J. Gollub, H. Jin, D. Botstein, J. M. Cherry, and G. Sherlock. GO::TermFinder—open source software for accessing Gene Ontology information and finding significantly enriched Gene Ontology terms associated with a list of genes. Bioinformatics, 20(18): 3710–3715, 2004.

9. S. Chib and E. Greenberg. Understanding the Metropolis-Hastings Algorithm. The American Statistician, 49(4):327–335, 1995.

10. V. Chvatal. A Greedy Heuristic for the Set-Covering Problem. Mathematics of Operations Research, 4(3): 233–235, 1979.

11. L. Geistlinger, G. Csaba, M. Santarelli, M. Ramos, L. Schiffer, N. Turaga, C. Law, S. Davis, V. Carey, M. Morgan, R. Zimmer, and L. Waldron. Toward a gold standard for benchmarking gene set enrichment analysis. Briefings in Bioinformatics, 22(1):545–556, 2021.

12. A. Gelman, J. B. Carlin, H. S. Stern, and D. B. Rubin. Bayesian Data Analysis. Chapman and Hall/CRC, 2003.

13. S. Grossmann, S. Bauer, P. N. Robinson, and M. Vingron. Improved detection of overrepresentation of Gene-Ontology annotations with parent–child analysis. Bioinformatics, 23 (22):3024–3031, 2007.

14. S. Holm. A Simple Sequentially Rejective Multiple Test Procedure. Scandinavian Journal of Statistics, 6(2):65–70, 1979.

15. A. Hukku, C. Quick, F. Luca, R. Pique-Regi, and X. Wen. BAGSE: A Bayesian hierarchical model approach for gene set enrichment analysis. Bioinformatics, 36(6):1689–1695, 2020.

16. S. Isci, C. Ozturk, J. Jones, and H. H. Otu. Pathway analysis of high-throughput biological data within a Bayesian network framework. Bioinformatics, 27(12):1667–1674, 2011.

17. P. Khatri, M. Sirota, and A. J. Butte. Ten Years of Pathway Analysis: Current Approaches and Outstanding Challenges. PLoS Computational Biology, 8(2):e1002375, 2012.

18. W. S. Noble. How does multiple testing correction work? Nature Biotechnology, 27(12):1135–1137, 2009.

19. I. Rivals, L. Personnaz, L. Taing, and M.-C. Potier. Enrichment or depletion of a GO category within a class of genes: Which test? Bioinformatics, 23(4):401–407, 2007.

20. P. N. Robinson and S. Bauer. Introduction to Bio-Ontologies. Chapman and Hall/CRC, 2011.

21. P. N. Robinson, A. Wollstein, U. B öhme, and B. Beattie. Ontologizing gene-expression microarray data: Characterizing clusters with Gene Ontology. Bioinformatics, 20(6):979–981, 2004.

22. A. Subramanian, P. Tamayo, V. K. Mootha, S. Mukherjee, B. L. Ebert, M. A. Gillette, A. Paulovich, S. L. Pomeroy, T. R. Golub, E. S. Lander, and J. P. Mesirov. Gene set enrichment analysis: A knowledge-based approach for interpreting genome-wide expression profiles. Proceedings of the National Academy of Sciences, 102(43):15545–15550, 2005.

23. R. Z. Vêncio, T. Koide, S. L. Gomes, and C. A. De B Pereira. BayGO: Bayesian analysis of ontology term enrichment in microarray data. BMC Bioinformatics, 7(1):86, 2006.

